# Cost-effectiveness of telephone-based weight loss support for patients with knee osteoarthritis: Results of an economic evaluation alongside a pragmatic randomised controlled trial

**DOI:** 10.1101/284588

**Authors:** KM O’Brien, JM van Dongen, A Williams, SJ Kamper, J Wiggers, RK Hodder, E Campbell, EK Robson, R Haskins, C Rissel, CM Williams

**Author notes:** Corresponding author: Kate M O’Brien.

## Abstract

**Background:** The prevalence of knee osteoarthritis is increasing worldwide. Obesity is an important modifiable risk factor for both the incidence and progression of knee osteoarthritis. Consequently, international guidelines recommend patients with knee osteoarthritis who are overweight receive support to lose weight. However, few overweight patients with knee osteoarthritis receive care to support weight loss. Telephone-based interventions are one potential solution to provide scalable care to the many patients with knee osteoarthritis. The objective of this study is to evaluate whether referral to a telephone-based weight management and healthy lifestyle service for patients with knee osteoarthritis, who are overweight or obese, is cost-effective from a societal perspective compared to usual care.

**Methods:** A cost-effectiveness analysis was undertaken alongside a pragmatic randomised controlled trial. We randomised 120 patients with knee osteoarthritis to an intervention or usual care control group in a 1:1 ratio. Participants in the intervention group received a referral to an existing non-disease specific 6-month telephone-based weight management and healthy lifestyle service. For this economic evaluation, the primary outcome was quality-adjusted life years (QALYs) and secondary outcomes included pain intensity, disability, weight, and body mass index (BMI). Costs included intervention costs, healthcare utilisation costs (healthcare services and medication use) and absenteeism costs due to knee pain. The primary cost-effectiveness analysis was performed from the societal perspective.

**Results:** Mean cost differences between groups (intervention minus control) were, $454 (95%CI: −2735 to 4206) for healthcare costs, $−36, (95%CI: −73 to 2) for medication costs, and $−13 (95%CI: −225 to 235) for absenteeism costs. The total mean difference in societal costs was $1022 (95%CI: −2201 to 4771). For all outcomes, the probability of the intervention being cost-effective compared with usual care was less than 0.33 at all willingness-to-pay values.

**Conclusions:** From a societal perspective, telephone-based weight loss support, provided using an existing non-disease specific 6-month weight management and healthy lifestyle service was not cost-effective in comparison with usual care for overweight and obese patients with knee osteoarthritis for QALYs, pain intensity, disability, weight, and BMI.

**Trial Registration number** ACTRN12615000490572

## Background

Osteoarthritis is one of the fastest growing chronic health problems worldwide.^1,2^ According to the 2015 Global Burden of Disease Study, osteoarthritis accounted for 3.9% of years lived with disability worldwide in 2015, up from 2.5% in 2010, and was the 13^th^ highest contributor to global disability.^1,2^ Knee osteoarthritis consistently accounts for approximately 85% of the burden attributable to osteoarthritis.^1,2^ Osteoarthritis also imposes a significant economic burden, with the total annual costs estimated to be $A8.6 billion in Australia,^3^ £20.9 billion in the UK,^4^ and $US142.1 billion in the US.^5^ The majority of these costs are attributable to ambulatory and inpatient care, including surgery and lost work productivity.^3,4^

Excess weight is an important modifiable risk factor for the onset and progression of knee osteoarthritis,^6^ and there is strong evidence that weight loss interventions reduce pain and disability in overweight patients with knee osteoarthritis.^7,8^ Consequently, international clinical practice guidelines recommend all patients with knee osteoarthritis who are overweight receive support to lose weight.^9–11^ Typically, these treatments are delivered using clinical face-to-face models of care.^12^ While such clinical models produce moderate effects on weight loss, pain, and physical function,^7,8^ only 22% of patients with knee osteoarthritis report receiving weight loss care,^13^ possibly due to limitations in service delivery and patient access to care. Arguably more scalable delivery options, using remotely delivered approaches, such as telephone-based support, can maximise the reach of weight loss care and are more cost-effective to support weight loss in this patient group. While telephone-based behavioural interventions targeting weight loss are used routinely in the general populations, the cost-effectiveness of referring patients with knee osteoarthritis to these is unknown.

Given the scarce resources in healthcare, policy-makers are increasingly requiring evidence of economic value for healthcare interventions to make informed decisions about how to allocate resources.^14^ Therefore, undertaking economic evaluations of knee osteoarthritis management approaches is important. Recently, we conducted a randomised controlled trial (RCT) using an existing non-disease specific telephone-based weight management and healthy lifestyle service for patients with knee osteoarthritis who are overweight or obese.^15^ The primary aim of the intervention was to reduce pain intensity, by reducing weight. The trial showed no between-group differences in pain intensity, nor weight.^15^ Despite the absence of clinical benefit, conducting a cost-effectiveness analysis is recommended.^14^ The cost-effectiveness analysis considers the joint distribution of differences in cost and effect and can show that an intervention is cost-effective when neither cost nor effect differences are individually significant.^14^ Cost-effectiveness analyses estimate the cost (saved or spent) per unit of effect gained. Such estimates can assist decision makers in prioritising interventions to determine how to best allocate limited funds. The purpose of the current study is to evaluate the cost-effectiveness of the aforementioned RCT, compared to usual care.

Economic analyses can be performed from various perspectives including the societal, and healthcare perspectives.^16^ The societal perspective includes all costs regardless of who pays. This frequently incorporates direct costs; intervention costs, plus costs of care unrelated to the intervention (i.e. healthcare services and medication costs), and the indirect costs; absence from work and impact on productivity.^16,17^ In contrast, the healthcare perspective only includes direct costs i.e. intervention costs and the costs of other care.^16^ In this study, the primary analysis was conducted from a societal perspective and a secondary analysis was conducted from the healthcare perspective.

## Methods

### Study participants and design

The economic evaluation was conducted alongside a two-arm pragmatic parallel group RCT, which was part of cohort multiple RCT.^18^ Full details of the study design has been described in the paper presenting the clinical results of the trial^15^ and in the study protocol.^19,20^ The trial was prospectively registered (ACTRN12615000490572). Ethical approval was obtained from The Hunter New England Health Human Research Ethics Committee (13/12/11/5.18) and the University of Newcastle Human Research Ethics Committee (H-2015-0043).

Patients with knee osteoarthritis who were on a waiting list for an outpatient orthopaedic consultation at the John Hunter Hospital in NSW, Australia, were invited to participate. Patients were assessed for eligibility during a telephone assessment. Eligible patients were randomised to study conditions: i) offered the intervention (intervention group), or ii) usual care control group.

Inclusion criteria were: primary complaint of pain due to knee osteoarthritis lasting longer than 3 months; 18 years or older; overweight or obese (body mass index (BMI) ≥27kg/m^2^ and <40kg/m^2^); average knee pain intensity ≥3 out of 10 on a 0-10 numeric rating scale over the past week, or moderate level of interference in activities of daily living (adaptation of item 8 of SF36); and access to a telephone. Exclusion criteria were: known or suspected serious pathology as the underlying cause of their knee pain (e.g. fracture; cancer, inflammatory arthritis; gout; or infection); previous obesity surgery; currently participating in any prescribed or commercial weight loss program; knee surgery in the last 6 months or planned surgery in the next 6 months; unable to comply with the study protocol that requires them to adapt meals or exercise due to non-independent living arrangements; medical or physical impairment precluding safe participation in exercise such as uncontrolled hypertension; and unable to speak or read English sufficiently to complete study procedures.

### Interventions

The intervention included two components. First, brief advice and education about the benefits of weight loss and physical activity for knee osteoarthritis were provided over the telephone immediately after randomisation. Second, intervention participants were informed about the NSW Get Healthy Information and Coaching Service (GHS) (www.gethealthynsw.com.au),^21^ and referred to the service for weight loss support. The GHS is an existing non-disease specific telephone-based health coaching service funded by the NSW Government to support adults to make sustained healthy lifestyle improvements. Targets include diet, physical activity and achieving a healthy weight, and where appropriate, referral to smoking cessation services.^21^ The GHS involves 10 individually tailored coaching calls, based on national dietary and physical activity guidelines,^22,23^ delivered over a 6-month period by university qualified health professionals.^21^ Participants in the intervention group remained on the waiting list for orthopaedic consultation.

Participants in the control group remained on the ‘usual care pathway’ (i.e. on the waiting list to have an orthopaedic consultation and could progress to consultation if scheduled or surgery if recommended by the orthopaedic department) and took part in data collection during the study period. No other active intervention was provided as part of the study, however; no restrictions were placed upon the use of other health services during the study period. Control participants were informed that a face-to-face clinical appointment was available in 6 months.

### Measures

For this economic evaluation, the primary outcome was quality-adjusted life years (QALYs). Secondary outcomes included pain intensity, disability, weight, and BMI. We measured cost in terms of intervention costs, healthcare utilisation costs (healthcare services and medication use) and work absenteeism costs due to knee pain. The primary analysis was conducted from the societal perspective, which includes all these cost categories. The secondary analysis was conducted from the healthcare perspective, which excluded absenteeism costs.

### Outcomes

All outcomes were assessed by self-reported questionnaires at baseline, six weeks and 26 weeks. The baseline questionnaire was completed by telephone. Week 6 and week 26 surveys were completed via telephone or mailed in the post as per participant preference. Health-related quality of life was assessed using the 12-item Short Form Health Survey version 2 (SF-12.v2).^24^ The participants’ SF-6D^25^ health states were translated into utility scores using the British tariff.^26^ QALYs were calculated by multiplying the participants’ utility scores by their time spent in a health state using linear interpolation between measurement points. Knee pain intensity was assessed using a 0-10 point NRS. Participants were asked to rate their “average knee pain intensity over the past week”, where 0 represents ‘no pain’ and 10 represents ‘the worst possible pain’.^27^ Disability was assessed using the Western Ontario and McMaster Universities Osteoarthritis Index (WOMAC).^28^ The total WOMAC score ranges from 0 to 96, with higher scores indicating greater disability. Weight (kg) was assessed via participant self-report and BMI was calculated as weight/height squared (kg/m^2^)^29^ using self-reported weight and height.

### Cost measures

All costs were converted to Australian dollars in 2016 using consumer price indices.^30^ Discounting of costs was not necessary due to the 26-week follow-up.^16^

Intervention costs were micro-costed and included the cost to provide the brief advice as well as GHS coaching calls. The cost to provide the brief advice was estimated from the development and operational costs of the call and the interviewer wages for the estimated average time (5 minutes) taken to provide the brief advice. The cost to provide the GHS coaching calls was provided by the GHS^31^ and multiplied by the number of calls each participant received. The number of health coaching calls received was reported directly by the GHS.

Healthcare utilisation costs included any healthcare services or medications used for knee pain (independent of the intervention costs) and were calculated from a patient reported healthcare utilisation inventory. Participants were asked to recall any healthcare services (the type of services and number of sessions) and medications used for their knee pain during the past six weeks, at the six and 26 weeks follow-up as part of the participant surveys. Healthcare services were valued using Australian standard costs and, if unavailable, prices were according to professional organisations.^32–34^ Medications were valued using unit prices from the Australian pharmaceutical benefits scheme^35^ and, if unavailable, prices were obtained from online Australian pharmacy websites. To gain an estimate of the cost of healthcare utilisation over the entire 26-week period, the average of the week six and week 26 costs per patient was extrapolated, assuming linearity.

Absenteeism was assessed by asking participants to recall the total number of sickness absence days from paid work due to knee pain during the past six weeks, at the six and 26 weeks follow-up as part of the participant surveys. Absenteeism costs were estimated using the ‘Human Capital Approach’^16^ and were calculated per participant by multiplying their total number of sickness absence days off by the national average hourly income for their gender and age according to the Australian Bureau of Statistics.^27,34^ To gain an estimate of the cost of absenteeism over the entire 26-week period, the average of the week six and week 26 costs per patient was extrapolated, assuming linearity.

### Statistical analysis

All outcomes and cost measures were analysed under the intention-to-treat principle (i.e. analyses were based on initial group assignment and missing data were imputed). Means and proportions of baseline characteristics were compared between the intervention and control group participants to assess comparability of the groups. Missing data for all outcomes and cost measures were imputed using Multiple Imputation by Chained Equations, stratified by treatment group.^36^ Data were assumed missing at random. Thirty different datasets were created in order for the loss-of-efficiency to be below the recommended 5%.^36^ These separate datasets were analysed as indicated below, after which pooled estimates from all imputed datasets were calculated using Rubin’s rules.^37^

Mean cost differences between study groups were calculated for total and disaggregated costs. The cost measures were adjusted for the confounders of baseline knee pain intensity, baseline duration of knee pain, baseline BMI and number of days on the waiting list for orthopaedic consultation because the addition of these confounders to the regression model changed the cost differences by more than 10%. Seemingly unrelated regression analyses were performed to estimate total cost differences (i.e. ΔC) and effect differences for all outcomes (i.e. ΔE), adjusted for baseline values as well as other potential baseline prognostic factors (knee pain intensity, duration of knee pain, BMI and number of days on the waiting list for orthopaedic consultation, obtained from hospital records). An advantage of seemingly unrelated regression is that two regression equations (i.e., one for ΔC and one ΔE) are modelled simultaneously so that their possible correlation can be accounted for.^38^

Incremental cost-effectiveness ratios (ICERs) were calculated by dividing the adjusted difference in total costs between both groups by the adjusted difference in outcomes (i.e. ΔC/ΔE). Uncertainty surrounding the ICERs and 95% confidence intervals (95%CIs) around cost differences were estimated using bias-corrected and accelerated bootstrapping (5000 replications). Uncertainty surrounding the ICERs was graphically illustrated by plotting bootstrapped incremental cost-effect pairs (CE-pairs) on cost-effectiveness planes (CE-planes).^16^ A summary measure of the joint uncertainty of costs and outcomes was provided using cost-effectiveness acceptability curves, which provide an indication of the intervention’s probability of being cost-effective in comparison with usual care at different values of willingness-to-pay (i.e., the maximum amount of money decision-makers are willing to pay per unit of effect).^16^ Data were analysed in STATA (V13, Stata Corp).

### Sensitivity analysis

To assess the robustness of results, we performed a per-protocol sensitivity analysis from the societal perspective that included only participants that completed at least six telephone GHS coaching calls in the intervention group.

### Secondary analysis: healthcare perspective

A secondary analysis was performed from the healthcare perspective (i.e. excluding absenteeism costs).

## Results

A total of 120 patients were randomised into the study (Figure 1). Baseline participant characteristics at baseline were similar between groups (Table 1). At 26 weeks complete data were available for between 70% and 82% of participants (QALYs 70%, pain intensity 82%, disability 79%, weight 81%, BMI 81%). For cost data, 48% of participants had complete data at 26 weeks. As a consequence, 18%-30% of outcome data and 52% of cost data were imputed.

**Table 1.** Baseline characteristics of the study population*

| Characteristics | Intervention group<br>*n*=59 | Control group<br>*n*=60 |
| --- | --- | --- |
| Age (years) | 63.0 (11.1) | 60.2 (13.9) |
| Gender (male), *n* (%) | 20 (34) | 25 (42) |
| Australian and/or Torres Strait Islander, *n* (%) | 5 (9) | 2 (3) |
| Employment status, *n* (%) |  |  |
| Employed | 12 (20) | 14 (23) |
| Unemployed | 7 (12) | 8 (13) |
| Retired | 31 (53) | 28 (47) |
| Can't work (health reasons) | 9 (15) | 10 (17) |
| Country of origin (Australia), *n* (%) | 54 (92) | 51 (85) |
| Highest level of education (>High school), *n* (%) | 11 (19) | 17 (28) |
| Private health insurance, *n* (%) | 1 (2) | 5 (8) |
| Pain intensity (NRS, range: 0-10) | 6.9 (1.8) | 6.8 (2.0) |
| Pain duration (years) | 9.6 (10.6) | 6.7 (8.5) |
| Disability (WOMAC, range: 0-96) | 47.9 (17.4) | 48.6 (16.5) |
| Weight (kg) | 93.3 (12.9) | 89.5 (13.5) |
| BMI (kg/m<sup>2</sup>) | 33.4 (3.4) | 32.1 (3.1) |
| Number of days waiting for orthopaedic consultation, median (IQR) | 379.0 (279.0-507.0) | 390.0 (313.0-532.0) |
| Utility score | 0.6 (0.1) | 0.7 (0.1) |

NRS=numeric rating scale, WOMAC=Western Ontario and McMaster Universities Osteoarthritis Index, BMI=Body Mass Index, IQR=interquartile range
\*Data presented as mean (SD) unless otherwise indicated.

**Figure 1.**
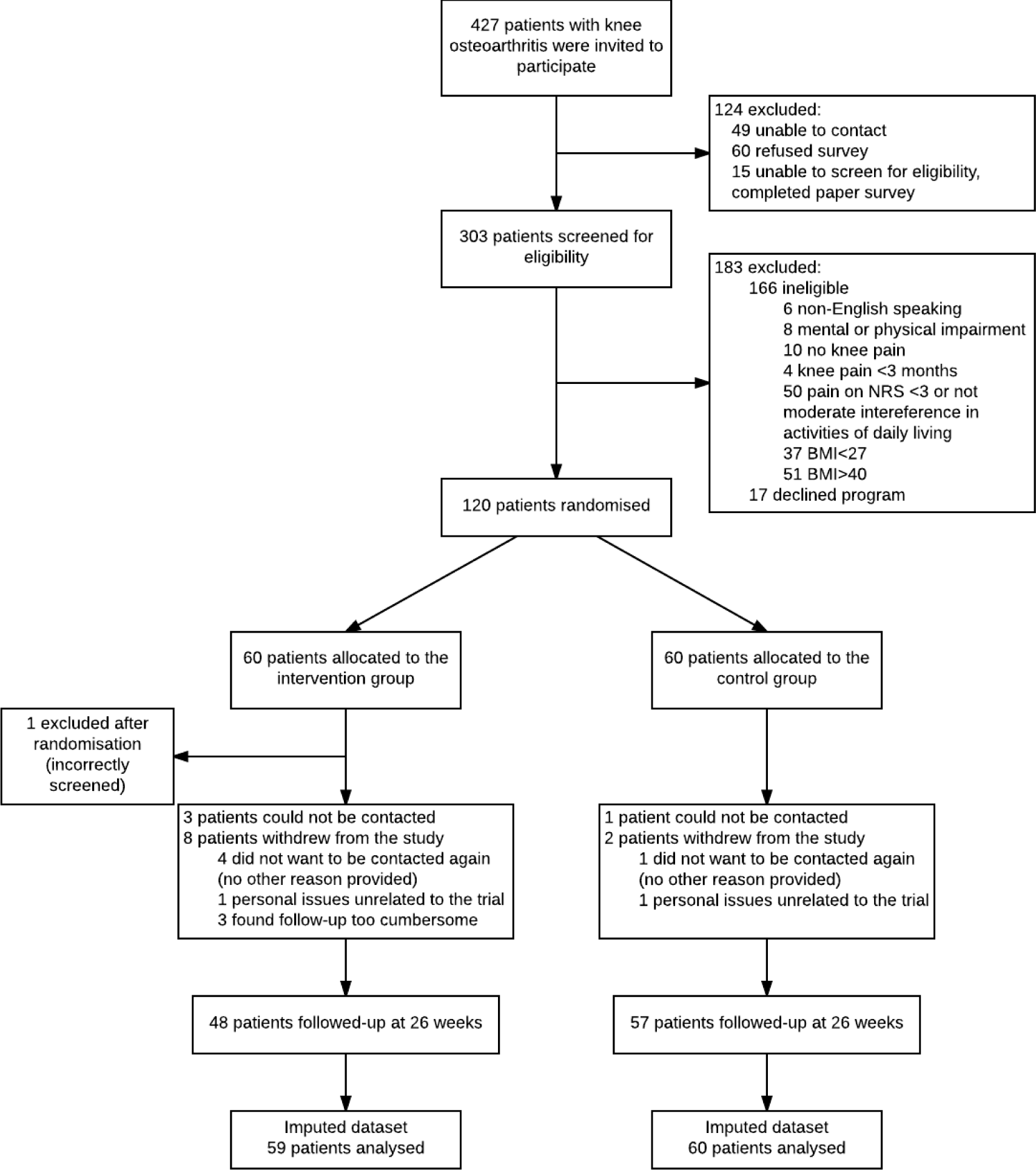
Flow diagram of trial participants

### Outcomes

No differences were found between the intervention and control group for QALYs (MD 0.00, 95%CI: −0.02 to 0.03), pain intensity (MD 0.58, 95%CI: −0.57 to 1.74), disability (MD −0.01, 95%CI: −8.00 to 7.99), weight (MD 0.36, 95%CI: −2.87 to 3.59), and BMI (MD 0.27, 95%CI: −0.93 to 1.46) (Table 2).

### Costs

Of the intervention group participants, the average number of GHS coaching calls to intervention participants was 4.7 (SD 4.6). The mean intervention costs were $622 (SE 80) per participant (Table 3).

From the societal perspective, mean cost differences between groups (intervention minus control) were, $454 (95%CI: −2735 to 4206) for healthcare costs, $−36, (95%CI: −73 to 2) for medication costs, and $−13 (95%CI: −225 to 235) for absenteeism costs. The total mean difference in societal costs was $1022 (95%CI: −2201 to 4771) (Table 3). From the healthcare perspective, total mean difference between groups (intervention minus control) was $876 (95%CI: −2419 to 4575).

### Cost-effectiveness

For QALYs, the majority of the incremental CE-pairs were located in the northeast quadrant, indicating the intervention was on average more costly and more effective than usual care (Figure 2 (1a)). The ICER for QALYs was 483,453, indicating that one QALY gained in the intervention group was associated with a societal cost of $483,453 as compared with the control group (Table 2). This large ICER is due to the very small effect on QALYs (MD 0.00, 95%CI: −0.02 to 0.03). The cost-effectiveness acceptability curve for QALYs is presented in Figure 2 (2a).

**Table 2.** Differences in pooled mean costs and effects (95% Confidence intervals), incremental cost-effectiveness ratios, and the distribution of incremental cost-effect pairs around the quadrants of the cost-effectiveness planes

| Analysis |  | Sample size |  | Outcomes | ΔC (95% CI) | ΔE (95% CI) | ICER | Distribution CE-plane (%) |  |  |  |
| --- | --- | --- | --- | --- | --- | --- | --- | --- | --- | --- | --- |
|  |  | Int | Cont |  | AUD | Points | AUD/point | NE<sup>a</sup> | SE<sup>b</sup> | SW<sup>c</sup> | NW<sup>d</sup> |
| Primary analysis | Societal perspective | 59 | 60 | QALYs | 1022 (-2217 to 4745) | 0.00 (-0.02 to 0.03) | 483453 | 37.6 | 18.7 | 14.2 | 29.4 |
|  |  | 59 | 60 | Pain intensity | 1022 (-2227 to 4760) | 0.58 (-0.57 to 1.74) | 1759 | 9.5 | 5.3 | 27.7 | 57.5 |
|  |  | 59 | 60 | Disability | 1022 (-2205 to 4725) | -0.01 (-8.00 to 7.99) | -178860 | 33.3 | 16.1 | 16.9 | 33.7 |
|  |  | 59 | 60 | Weight | 1022 (-2218 to 4717) | 0.36 (-2.87 to 3.59) | 2851 | 28.5 | 13.4 | 19.3 | 38.7 |
|  |  | 59 | 60 | BMI | 1022 (-2225 to 4750) | 0.27 (-0.93 to 1.46) | 3850 | 23.7 | 10.9 | 22.0 | 43.4 |
| Sensitivity analysis | Per protocol | 59 | 60 | QALYs | 192 (-3692 to 4303) | 0.01 (-0.03 to 0.04) | 22985 | 37.0 | 31.6 | 14.1 | 17.3 |
|  |  | 59 | 60 | Pain intensity | 192 (-3710 to 4303) | 0.56 (-0.98 to 2.10) | 340 | 12.6 | 10.7 | 35.0 | 41.7 |
|  |  | 59 | 60 | Disability | 192 (-3684 to 4316) | -0.04 (-11.40 to 11.48) | -5347 | 27.9 | 21.6 | 24.1 | 26.4 |
|  |  | 59 | 60 | Weight | 192 (-3678 to 4287) | 1.32 (-4.12 to 6.76) | 146 | 17.2 | 14.8 | 30.9 | 37.1 |
|  |  | 59 | 60 | BMI | 192 (-3658 to 4317) | 0.74 (-1.27 to 2.74) | 261 | 13.3 | 11.1 | 34.5 | 41.1 |
| Secondary analysis | Healthcare perspective | 59 | 60 | QALYs | 758 (-2439 to 4439) | 0.00 (-0.02 to 0.03) | 358615 | 34.9 | 21.3 | 16.4 | 27.4 |
|  |  | 59 | 60 | Pain intensity | 758 (-2448 to 4456) | 0.58 (-0.58 to 1.74) | 1305 | 8.9 | 6.1 | 31.8 | 53.2 |
|  |  | 59 | 60 | Disability | 758 (-2460 to 4409) | -0.00 (-8.00 to 8.00) | -164545 | 30.5 | 18.8 | 19.1 | 31.6 |
|  |  | 59 | 60 | Weight | 758 (-2434 to 4466) | 0.36 (-2.88 to 3.60) | 2115 | 26.3 | 15.7 | 22.2 | 35.8 |
|  |  | 59 | 60 | BMI | 758 (-2431 to 4442) | 0.27 (-0.93 to 1.46) | 2856 | 21.8 | 12.4 | 25.3 | 40.5 |

Int=Intervention, Cont=Control, CI=confidence interval, C=costs, E=effects, ICER=incremental cost-effectiveness ratio, CE-plane=cost-effectiveness plane, BMI=Body mass index, QALYs=quality-adjusted life years
Note: costs are expressed in 2016 Australian Dollars
<sup>a</sup> The northeast (NE) quadrant of the CE plane, indicating that the intervention is more effective and more costly than control
<sup>b</sup> The southeast (SE) quadrant of the CE plane, indicating that the intervention is more effective and less costly than control
<sup>c</sup> The southwest (SW) quadrant of the CE plane, indicating that the intervention is less effective and less costly than control
<sup>d</sup> The northwest (NW) quadrant of the CE plane, indicating that the intervention is less effective and more costly than control

**Table 3.** Mean costs per participant in the intervention and control groups, and unadjusted and adjusted mean cost differences between study groups during the 6-month follow-up period (based on the imputed dataset)

|  | Intervention group<br>*n*=59 | Control group<br>*n*=60 | Unadjusted mean<br>cost difference | Adjusted mean cost<br>difference* |
| --- | --- | --- | --- | --- |
| Cost category | mean (SE) | mean (SE) | CI (95%) | CI (95%) |
| Intervention costs | 622 (80) | 0 (0) | 622 (480 to 792) | 618 (464 to 807) |
| Healthcare costs | 2728 (1747) | 2605 (1463) | 312 (-2728 to 3533) | 454 (-2735 to 4206) |
| Medication costs | 108 (24) | 139 (22) | -35 (-71 to 1) | -36 (-73 to 2) |
| Absenteeism costs | 174 (124) | 160 (72) | 17 (-170 to 236) | -13 (-225 to 235) |
| **Total** | **3632 (1781)** | **2904 (1478)** | **916 (-2163 to 4135)** | **1022 (-2201 to 4771)** |

SE=standard error, CI=confidence interval
Note: costs are expressed in 2016 Australian Dollars
Negative difference values indicate control group costs greater than intervention
\*Mean cost difference (intervention minus control) adjusted for the baseline variables: knee pain intensity, duration of knee pain (years), BMI, number of days on the waiting list for orthopaedic consultation.

**Figure 2.**
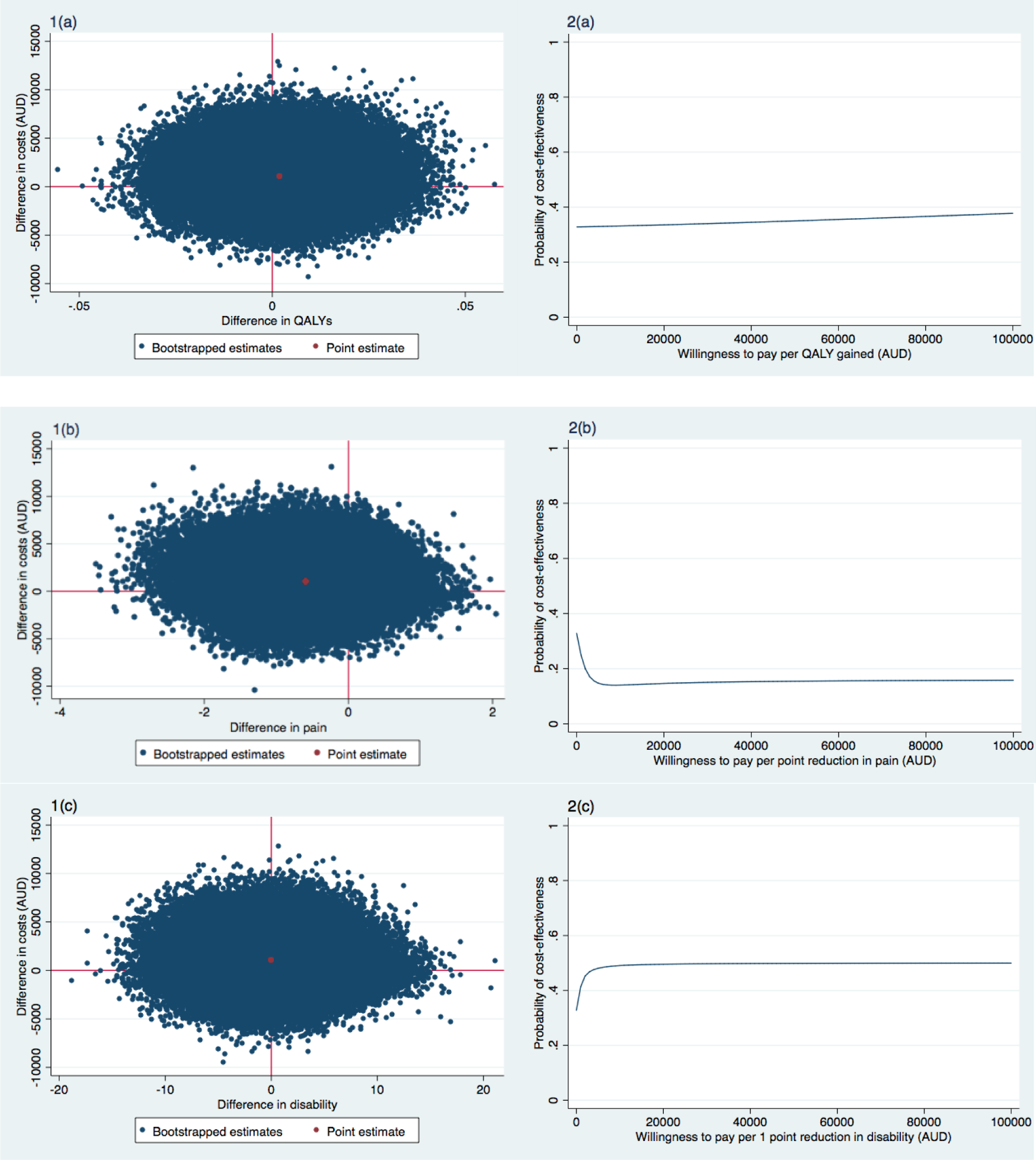
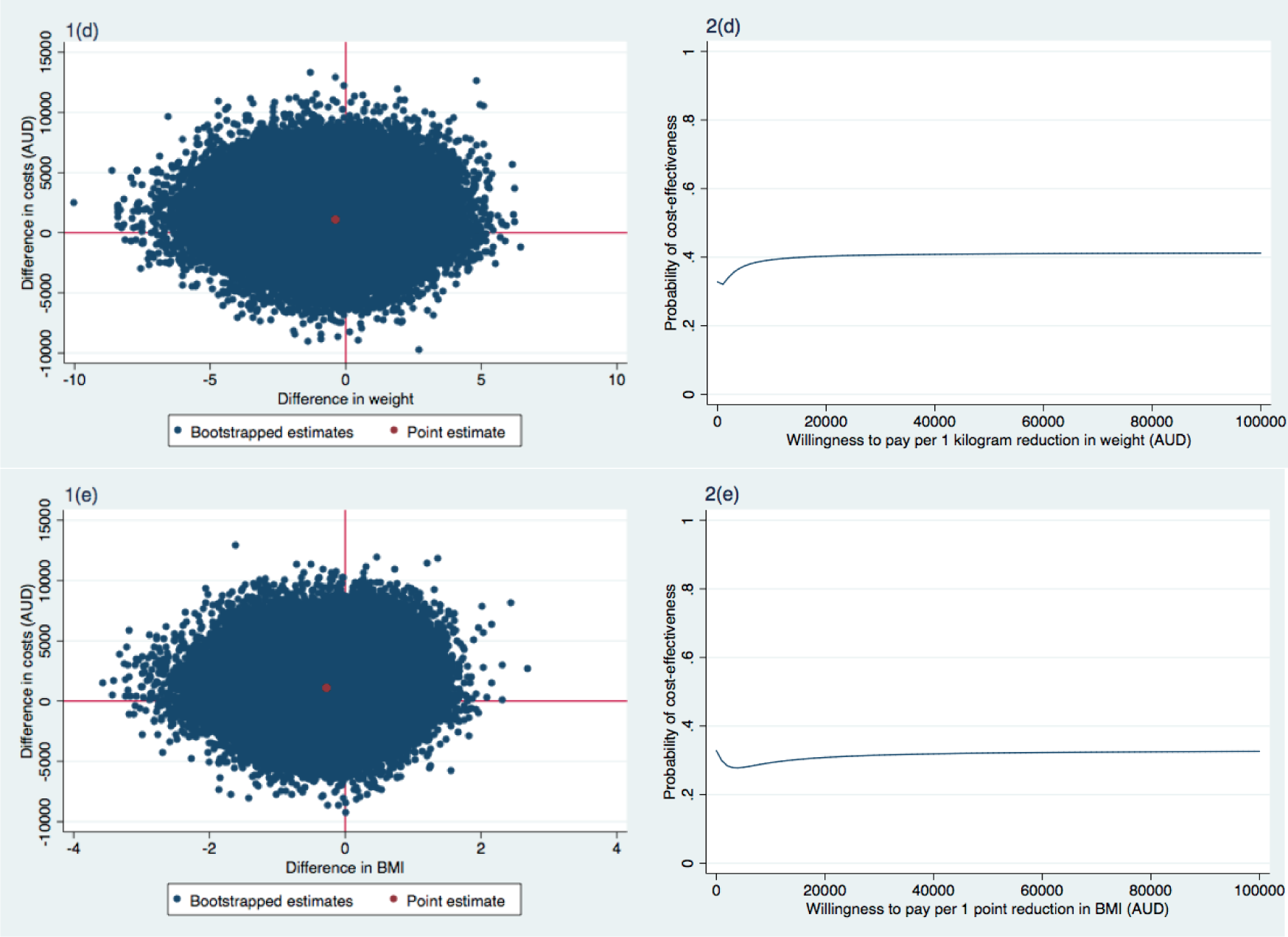
Cost-effectiveness planes indicating the uncertainty around the incremental cost-effectiveness ratios (1) and cost-effectiveness acceptability curves indicating the probability of cost-effectiveness for different values ($) of willingness-to-pay per unit of effect gained (2) for QALY (a), pain intensity (b), disability (c), Weight (d), and BMI (e) (based on the imputed dataset)

For the secondary outcomes, the majority of incremental CE-pairs were located in the northwest quadrant (Table 2, Figure 2 (1b-1e)), indicating that the intervention on average achieved poorer outcomes at a higher cost compared to usual practice. As such these ICERs had to be interpreted as the cost per unit of effect lost. For pain intensity, an ICER of 1,759 indicated that the intervention costs $1,759 per point increase in pain intensity in comparison to the control group (Table 2, Figure 2 (1b)). ICERs for weight (2,851) and BMI (3,850) in a similar direction were found, which indicated the intervention costs $2,851 per one kilogram gained and $3,850 per one unit BMI increase (Table 2, Figure 2 (1d and 1e)). In contrast, the ICER for disability was −178,860, which indicated for every 1-point decrease in disability (i.e. improvement), the intervention costs $178,860 compared with the control group (Table 2, Figure 2 (1c)). This large ICER was due to the very small effect on disability (MD −0.01, 95%CI: −8.00 to 7.99). Cost-effectiveness acceptability curves for pain intensity, disability, weight, and BMI are presented in Figure 2 (2b-2e).

For all outcomes, the probability of the intervention being cost-effective in comparison to usual care was 0.33 at a willingness-to-pay of $0/unit of effect gained. For QALYs, disability, weight, and BMI the probability remained about the same irrespective of the willingness-to-pay (Figure 2 (2a, 2c-2e)). For pain intensity, this probability decreased with increasing values of willingness-to-pay (Figure 2 (2b)).

### Sensitivity analysis

Results of the sensitivity analysis can be found in Table 2. In brief, for QALYs, the probability of cost-effectiveness was 0.47 at a willingness-to-pay of $0 per QALY gained. For QALYs and disability, the probability of cost-effectiveness remained about the same irrespective of the willingness-to-pay. For pain intensity, weight and BMI the probability decreased for increasing values of willingness-to-pay.

### Secondary analysis: healthcare perspective

The ICER for QALY was 358,615 indicating that one QALY gained was associated with a cost of $358,615 compared with the control group (Table 2).

For pain intensity, the ICER was 1,305, indicating that a one-point increase in pain intensity was associated with a cost of $1,305, indicating that the intervention was on average more costly and less effective than usual practice (i.e. achieving poorer outcomes at a higher cost).

ICERs in similar directions were found for weight ($2,115 per one kilogram gained) and BMI ($2,856 per one-point increase) (Table 2). For disability, an ICER of −164,545 was found. This indicates that for every 1-point decrease (i.e. improvement) in disability, the intervention costs $164,545 (Table 2).

For all outcomes, the probability of the intervention being cost-effective in comparison to usual care was 0.37 at a willingness-to-pay of $0/unit of effect gained. For QALYs, disability, and weight the probability remained about the same irrespective of the willingness-to-pay. For pain intensity and BMI, this probability decreased with increasing values of willingness-to-pay.

## Discussion

We found that referral to a telephone-based weight management and healthy lifestyle service was not cost-effective from a societal perspective compared with usual care, for patients with knee osteoarthritis who are overweight or obese. Sensitivity analyses confirmed these results. The maximum probability of the intervention being cost-effective was low for all outcomes and perspectives, irrespective of the willingness-to-pay.

To our knowledge, there are no other economic evaluations of telephone-based interventions for patients with knee osteoarthritis, yet our study supports other studies showing limited evidence of cost-effectiveness for weight loss treatments for osteoarthritis. A systematic review of non-pharmacological and non-surgical interventions found from 11 cost-effectiveness studies, exercise programs appear to offer the most cost saving but there have been very few reports of the cost-effectiveness of conservative treatments compared to usual or minimal care controls.^39^ Of all four trials evaluating lifestyle programs (including weight loss), none were cost-effective.^39^ Importantly, the review reported less than 50% of studies to have acceptable methodological quality due to a combination of poor RCT quality, and poor quality economic analyses. Our study using a high-quality RCT and contemporary economic analytic methods shows that telephone support targeting weight loss behaviours, physical activity and diet was neither cost-saving nor cost-effective.

A more recent high-quality study assessed the cost-effectiveness of a 6-week multidisciplinary face-to-face treatment program compared with a telephone-based program for patients with generalised osteoarthritis.^40^ In this study, all patients received in-depth education with the overall goal to enhance self-management skills.^40^ Patients in the face-to-face group received six therapeutic groups session (2-4 hours each), whereas the telephone group received only two face-to-face group sessions (2-2.5 hours each) and four individual telephone contacts (15-30 mins each). The study found that from a societal perspective the face-to-face treatment was more likely to be cost-effective at 1-year follow-up.^40^ These results, together with the findings from our current study, suggest that telephone-based care for patients with osteoarthritis may not be a cost-effective management approach. Since many patients with osteoarthritis do not receive recommended treatments via clinical models of care,^13,41^ understanding why telephone-based interventions are reported to be as effective as face-to-face interventions but not cost-effective, is an important consideration to inform how best to provide care to this patient group.

An important strength of the present study is the pragmatic RCT design. The pragmatic design of the trial enabled us to evaluate the intervention under ‘real world’ circumstances. This facilitates the generalisability of the results and allows decision-makers to use these results to help guide future healthcare interventions. Another strength is the use of contemporary statistical methods. Multiple imputation was used to avoid problems of lost power and inefficiency associated with complete-case analyses. Seemingly unrelated regression analyses were used for analysing the cost and effect components of the cost-effectiveness analysis, allowing us to adjust for various potential confounders that may not be the same for costs and effects, while simultaneously accounting for the possible correlation between costs and effects. Bootstrapping techniques were used allowing for an estimation of uncertainty surrounding the cost-effectiveness estimates.

This study is not without limitations. Firstly, the rate of missing data at 26 weeks, between 18% and 30% for effect measures and 52% for cost data is high, however, not dissimilar to those in other economic evaluations.^42^ We used multiple imputation to account for the missing data, which is recommended over complete case analyses, despite this, results from this study should be treated with caution. Another limitation is that costs were obtained through self-reported questionnaires, which may have introduced recall bias, although the recall period was short (6 weeks). Lastly, presenteeism costs were not included, (i.e. reduced productivity while at work) which is known to be an issue reported by patients with chronic disabling pain.^43^

There is a need for more information about the cost-effectiveness of lifestyle interventions for osteoarthritis. Although this study indicates that the use of a generic non-disease specific telephone-based service is not cost-effective for overweight and obese patients with knee osteoarthritis, the current evidence suggests existing models of care delivery are unable to provide recommended care to the large number of patients with knee osteoarthritis.^13^ More research into how to provide scalable models of care that are cost-effective is needed.

Interestingly, the National Institute for Health and Clinical Excellence guidelines for the management of osteoarthritis^9^ refer to general obesity management guidelines for weight loss care for these patients,^44^ and not disease-specific models of care. Based on the results of our study, recommending general non-disease specific weight loss interventions may not be appropriate, or cost-effective for these patients. A key feature of our study and that of other osteoarthritis telephone interventions is that they only provide support over a relatively short period (six weeks to six months). However, other general weight loss programs occur over a much longer time frame. Better understanding about how the key ingredients for telephone services like dose and relevant components (e.g. exercise, weight loss, education) affect cost-effectiveness may provide more insight about the true value of telephone-based approaches for osteoarthritis.

## Conclusions

Our findings suggest that referral to a telephone-based weight management and healthy lifestyle service is not cost-effective compared with usual care for overweight and obese patients with knee osteoarthritis. These findings apply to QALYs, pain intensity, disability, weight, or BMI, from the societal and healthcare system perspectives.

## Ethics approval

Approval was obtained from the Hunter New England Health Human Research Ethics Committee (13/12/11/5.18) and the University of Newcastle Human Research Ethics Committee (H-2015-0043).

## Authors’ contributions

KO, CW, AW, JW, RH and EC were responsible for the concept and design of the study. CW and JW were responsible for obtaining trial funding. KO, JVD and CW drafted the manuscript, and all authors contributed to the interpretation of the data for the work and revision of the manuscript. All authors have read and approved the final manuscript. KO and CW take responsibility for the integrity of the work as a whole, from inception to finished article.

